# Microbiota may affect the tumor type but not overall tumor development in two models of heritable cancer

**DOI:** 10.1101/2023.10.11.561890

**Authors:** Jessica Spring, Sandeep Gurbuxani, Tatyana Golovkina

## Abstract

Microbial impact on tumorigenesis of heritable cancers proximal to the gut is well documented. Whether the microbiota influences cancers arising from inborn mutations at sites distal to the gut is undetermined. Using two models of heritable cancer, we found the microbiota to be inconsequential for tumor development. However, the type of tumor that develops may be influenced by the microbiota. This work furthers our understanding of the microbial impact on tumor development.

## Introduction

Cancer arises from sporadic or inborn mutations within oncogenes, tumor suppressor genes, or the regulatory regions that control their expression (1). Sporadic mutations occur randomly during normal cell division or emanate from radiation, chemical carcinogens, or viruses (2, 3). On the other hand, inborn mutations are inherited and can cause familial cancers (4, 5). The influence of the microbiota on carcinogenesis stemming from spontaneous mutations has been well documented by us and other researchers (6-8). Furthermore, the development and progression of hereditary colorectal cancer, often caused by mutations in tumor suppressor genes such as Adenomatous polyposis coli (APC), has been found to be influenced by the microbiota (6, 9). However, the potential role the microbiota plays in carcinogenesis of heritable cancers developing distal from the gut is not well understood.

A previous study of mice deficient in ataxia telangiectasia mutated (ATM) gene suggested the microbiota played a role in lymphomagenesis (10). ATM is activated by double stranded DNA breaks and by phosphorylating tumor-suppressor gene p53, induces cell cycle arrest or apoptosis (11). Mutations in this gene cause ataxia telangiectasia, also known as Louis-Bar syndrome in humans, an autosomal recessive disease that in addition to other symptoms, causes a higher incidence of lymphoid cancers (12). Yamamoto et al. found ATM-deficient mice exhibited increased lymphoma latency when housed in specific pathogen free (SPF) conditions and supplied with sterile food, water, and bedding in comparison to ATM-deficient mice housed in SPF conditions with non-autoclaved supplies (13). Furthermore, ATM-deficient mice gavaged with a restricted microbiota following antibiotic treatment had an increased lifespan compared to mice gavaged with a conventional microbiota after antibiotic treatment (10). While these data suggest an influence of the microbiota on hereditary cancer that forms in regions distal from the gut, more conclusive results would require the use of completely sterile germ-free (GF) mice.

To definitively determine whether the microbiota influences cancers arising from inborn mutations we used two animal models with predisposing genetic defects to analyze and compare the frequency of tumor development between SPF and GF settings.

## Materials and methods

### Mice

The following mice used in this study were bred and maintained at the animal facility of The University of Chicago. B6.129S2-*Trp53*^*tm1Tyj*^*/J* (B6.Trp53^-/-^) and FVB.Cg-Tg(Wnt1)1Hev/J (FBV.wnt1Tg) were purchased from The Jackson Laboratory.

### Monitoring GF isolator sterility

B6.Trp53^-/-^ and FBV.wnt1Tg mice were re-derived as germ-free (GF) at Taconic and housed at the gnotobiotic facility in sterile isolators at The University of Chicago. Assessment of GF isolator sterility was conducted as previously described (14). Briefly, fecal pellets collected from isolators weekly were frozen followed by DNA extraction using a bead-beating/phenol-chloroform extraction protocol. A single fecal pellet was placed in an autoclaved 2ml screw-cap tube containing 0.1mm zirconium beads along with 500µl of 2X buffer (filter sterilized 200mM NaCl, 200mM Tris, 20mM EDTA), 210µl of 20% SDS, and 500µl phenol:chloroform. The tube was bead beat on high for 2 minutes then centrifuged at 8,000 rpm at 4°C for 3 minutes. The aqueous phase was placed in a new Eppendorf tube and 500µl of phenol:chloroform was added. The tube was centrifuged at 13,000 rpm at 4°C for 3 minutes. The aqueous phase was again extracted, mixed with 40µl of 3M sodium acetate pH 7 and 400µl of -20°C isopropanol, and spun at 13,000 rpm at 4°C for 10 minutes. The supernatant was discarded and 500µl of - 20°C 80% ethanol was added. The sample was spun at 13,000 rpm at 4°C for 5 minutes. The supernatant was again dumped, and the sample was vacuum dried to 10 minutes. The pellet was resuspended in 1,000µl of sterile water and left overnight at 4°C. Primers that broadly hybridize to bacterial 16S rRNA gene sequences (5’GACGGGCGGTGWGTRCA3’ and 5’AGAGTTTGATCCTGGCTCAG3’) were used to amplify isolated DNA. Furthermore, microbiological cultures were inoculated with GF fecal pellets, positive control SPF fecal pellets, sterile saline (sham), and sterile culture medium (negative control). BHI, Nutrient, and Sabbaroud Broth tubes were inoculated and incubated at 37°C and 42°C aerobically and anaerobically. Cultures were monitored for five days until deemed negative.

### Histology

Visible tumors were excised from SPF and GF *Trp53*-deficient and Wnt1-transgenic mice and fixed in Telly’s fixative. Tumor type was determined based on morphologic assessment of hematoxylin and eosin stained 4-micron sections.

## Results

The first model tested was tumor suppressor p53 (encoded by *TRP53* in humans and *Trp53* in mice) deficiency. *TRP53* is crucial for enabling response to cell stress, such as DNA damage, by arresting cell-cycling or inducing apoptosis (15). Mutations within *TRP53* are the most frequently observed mutations in human cancers (16). In addition, germline mutations in *TRP53* are associated with Li-Fraumeni syndrome, a cancer with familial predisposition (17). Mice deficient in *Trp53* were rederived as germ-free and along with SPF *Trp53*-deficient mice, were monitored for tumor development. The frequency of tumor development in GF *Trp53*-deficient mice was very similar to that observed in SPF *Trp53*-deficient mice (Figure 1A). Furthermore, all SPF and the majority of GF *Trp53*-deficient mice developed lymphomas or sarcomas (Figures 1B and 1C) in agreement with previously published data (18). It is noteworthy that some GF *Trp53*-deficient mice developed carcinomas, a cancer type less frequently associated with this mouse model (19). This suggests the microbiota may preclude rather than promote this type of tumor.

**Figure 1.**
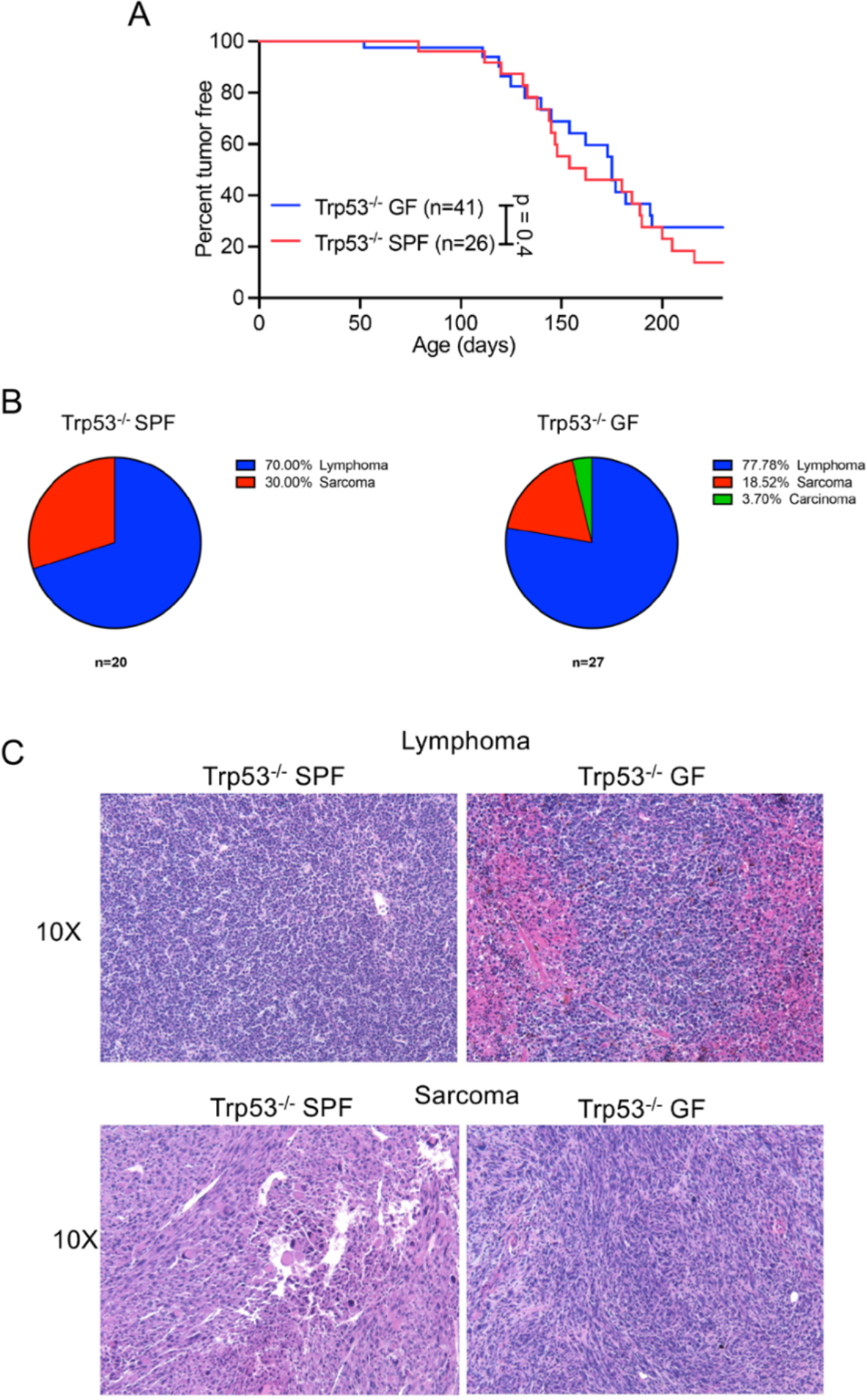
Microbiota does not impact the tumorigenesis but may influence tumor type in mice deficient in p53. **(A)** C57BL/6 Trp53^-/-^ SPF and GF mice were monitored for tumor development. **(B)** Proportion of SPF Trp53^-/-^ mice that develop various forms of tumors in SPF (left) and GF (right) setting. **(C)** Tumors were excised from the surrounding tissue, fixed, sectioned, and stained with hematoxylin-and eosin. Representative hematoxylin-and eosin stained lymphomas (top) and sarcomas (bottom) from Trp53^-/-^ SPF (left) and GF (right) mice. 10X magnification. n, number of mice used. *p* values calculated using Mantel-Cox test **(A)**.

To further understand the influence of the microbiota on tumor development stemming from genetic predisposition, we also tested Wnt1 transgenic mice, a model for mammary carcinoma development. Wnt1 canonically controls cell proliferation by increasing and stabilizing cytosolic β-catenin, which then translocates to the nucleus and facilitates the expression of genes including cell cycle regulators c-myc and cyclin D1 (20). Atypical Wnt1 expression under the mammary tumor virus promoter (MMTV-Wnt1) within Wnt1 transgenic mice induces development of mammary adenocarcinomas (20). GF MMTV-Wnt1 transgenic mice developed tumors at a similar rate to SPF MMTV-Wnt1 transgenic mice (Figure 2A). Wnt1 transgenic mice in both housing conditions primarily developed the archetypal tumor type for this model, adenocarcinomas (Figures 2B and 2C). However, a few GF MMTV-Wnt1 transgenic mice developed squamous cell carcinomas (Figure 2B), again suggesting the microbiota may be protective against some tumor types.

**Figure 2.**
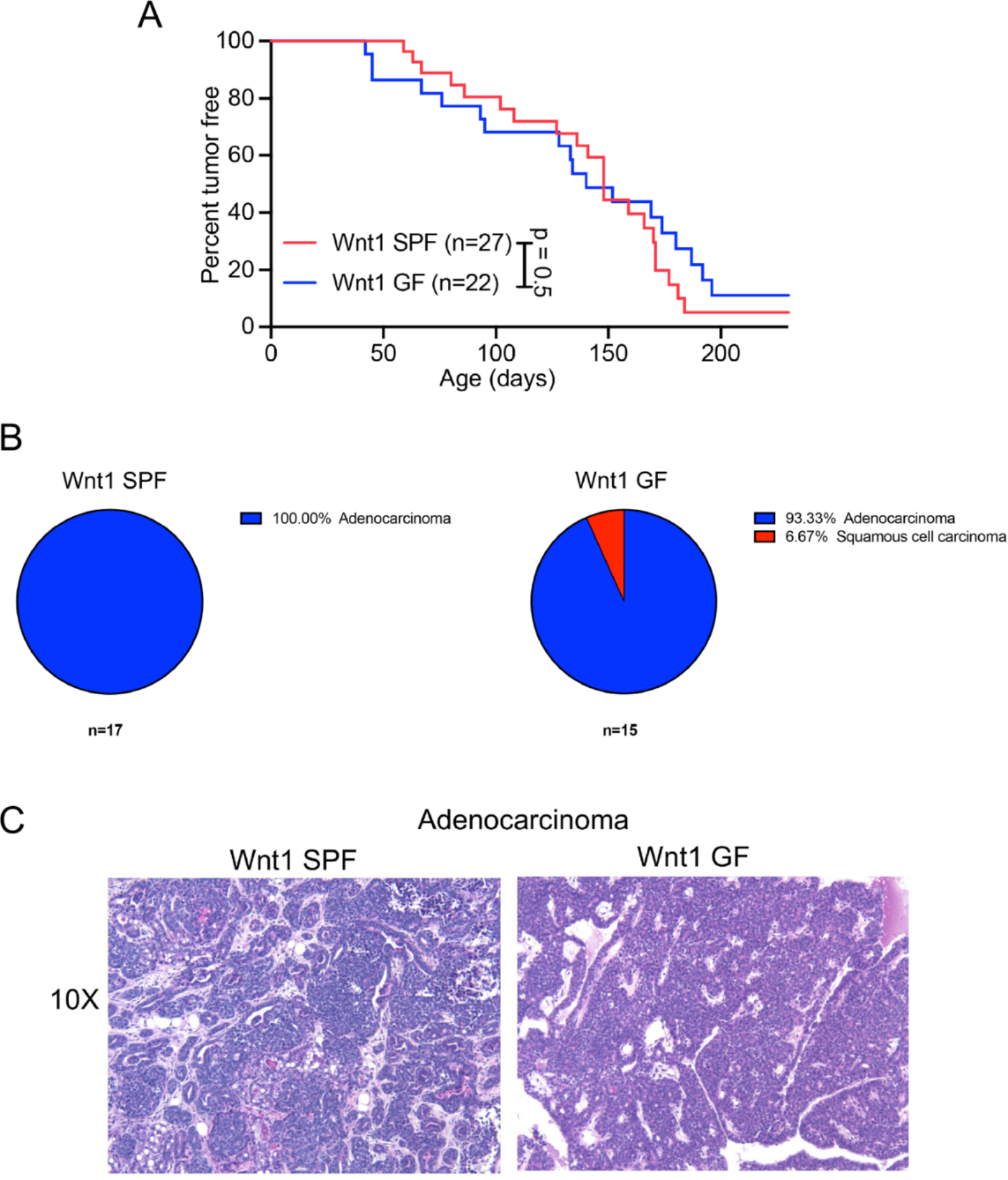
The microbiota does not influence tumor development but may impact tumor type in mice transgenic for Wnt1. **(A)** FVB Wnt1 transgenic SPF and GF mice were monitored for tumor development. **(B)** Proportion of Wnt1 transgenic mice that develop various types of tumors in SPF (left) and GF (right) setting. **(C)** Tumors were excised from the surrounding tissue, fixed, sectioned, and stained with hematoxylin-and eosin. Example hematoxylin-and eosin stained adenocarcinomas from Wnt1 transgenic SPF (left) and GF (right) mice. 10X magnification. n, number of mice used. *p* values calculated using Mantel-Cox test **(A)**.

## Discussion

In contrast to previous studies on spontaneous tumor development, including our study of leukemia induced by murine leukemia virus (MuLV) (6-8), the latency of hereditary lymphoma/sarcoma and mammary gland adenocarcinomas in *Trp53*^-/-^ and Wnt1 transgenic mice, respectively, were unaffected by the absence of the microbiota (Figures 1A and 2A). These data suggest that unlike spontaneous malignancies, the development of cancer in gut distant organs arising from genetic predisposition may not be influenced by the microbiota.

The discordance in these observations may be explained by the inherent differences between tumors that develop spontaneously, such as MuLV induced leukemia, and the tumors that develop in *Trp53*^-/-^ and Wnt1-transgenic mice. Spontaneous tumors, induced by MuLV (and those induced by other viruses or carcinogens), derive from a small percentage of unique, altered cells in which oncogenes are upregulated by provirus insertion. These cells express viral antigens or neoantigens which are foreign to the host. A robust immune response against these limited number of cells, as seen in GF MULV infected mice, renders these mice highly resistant to MuLV induced leukemia. In contrast, the presence of the microbiota in SPF MuLV infected mice suppresses the immune response enabling leukemia development.

Within models of heritable cancer, inborn mutations are ubiquitous in all cells of the body or particular tissue. As all cells have already acquired the first mutation necessary for tumor development, foreign antigens are likely absent at the initiation of tumorigenesis. As cells evolve and develop further mutations, few neoantigens may reach lymphoid organs, leading to immunological ignorance where the antigen is neglected by the immune system (21-23).

Additionally, tumor cells reaching the lymph node can utilize various mechanisms to prevent activation of the immune response and contribute to immunological ignorance (21, 23). As these mutated cells undergo tumorigenesis, the overwhelming and consistently present tumor antigens in lymphoid organs could lead to exhaustion whereby an initial activation and proliferation of T cells is followed by deletion of these specific T cells (21, 24). These processes alone could drive tumorigenesis regardless of the presence or absence of tumor influencing microbiota.

While the microbiota does not appear to contribute to the frequency of tumor development in these models of inheritable cancers, the microbiota may affect the histological type of tumors that grow. In both *Trp53*-deficient and Wnt-transgenic mice, the absence of microbiota led to a higher frequency of development of certain type of carcinomas (Figures 1B and 2B). Thus, it is possible that microbiota has a negative effect on the development of epithelial tumors.

## Author contributions

J.S. performed the experiments reported in the paper. S.G. examined histology of tumor sections. J.S. and T.G. wrote the manuscript. T.G. conceived the project.

## Acknowledgements

We thank members of the Golovkina lab for helpful discussions. We also thank GRAF at University of Chicago which enable gnotobiotic studies. This work was supported by NIH grants by R01CA232882 to T.G, J.S. was supported by T32 GM007183. This work was also supported by NIH/NIDDK Digestive Disease Research Core Center grant DK42086 and NIH grant P30CA014599.

